# EuMicrobedbLite: A lightweight genomic resource and analytic platform for draft oomycete genomes

**DOI:** 10.1101/084483

**Authors:** Arijit Panda, Diya Sen, Arup Ghosh, Akash Gupta, Malar C Mathu, Gyan Prakash Mishra, Deeksha Singh, Wenwu Ye, Brett M. Tyler, Sucheta Tripathy

**Affiliations:** Computational Genomics lab, Structural Biology and Bioinformatics Division, Council of Scientific and Industrial Research-Indian Institute of Chemical Biology, 4, Raja S.C Mullick Road, Jadavpur, Kolkata, 700032, India.; Center for Genome Research and Biocomputing and Department of Botany and Plant Pathology, Oregon State University, Corvallis, Oregon 97331-7303.; Department of Plant Pathology, Nanjing Agricultural University, Nanjing 210095, China

**Keywords:** Oomycetes, Comparative Genomics, Database, Genome Browser, Toolkit, Orthologous Genes

## Abstract

We have developed EuMicrobedbLite – A light weight comprehensive genome resource and sequence analysis platform for oomycete organisms. EuMicrobedbLite is a successor of the VBI Microbial Database (VMD) that was built using the Genome Unified Schema (GUS). In this version, the GUS schema has been greatly simplified with removal of many obsolete modules and redesign of others to incorporate contemporary data. Several dependencies such as perl object layers used for data loading in VMD have been replaced with independent light weight scripts. EumicrobedbLite now runs on a powerful annotation engine developed at our lab called “Genome Annotator Lite”. Currently this database has 26 publicly available genomes and 10 EST datasets of oomycete organisms. The browser page has dynamic tracks presenting comparative genomics analyses, coding and non-coding data, tRNA genes, repeats and EST alignments. In addition, we have defined 44,777 core conserved proteins from twelve oomycete organisms that form 2974 clusters. Synteny viewing is enabled by incorporation of the Genome Synteny Viewer (GSV) tool. The user interface has undergone major changes for ease of browsing. Queryable comparative genomics information, conserved orthologous genes and pathways are among the new key features updated in this database. The browser has been upgraded to enable user upload of GFF files for quick view of genome annotation comparisons. The toolkit page integrates the EMBOSS package and has a gene prediction tool. Annotations for the organisms are updated once every six months to ensure quality. The database resource is available at www.eumicrobedb.org.

**Abbreviations**
eumicrobedblite: GUS, VMD, GFF, GO, KEGG, KAAS, KOG, TMHMM

## Introduction

Many oomycetes are destructive pathogens against crop plants, animals and humans and pose a major threat to global food security (Dong et al., 2014, Pennisi, 2010). These pathogens were earlier believed to be fungi, mostly because of their morphology, but were later grouped under stramenopiles (Adhikari et al., 2013). The early progenitors of oomycetes have been proposed to be phototrophic brown algae that lost their ability to photosynthesize and became parasites (Tyler et al., 2006). While many pathogens and parasites have undergone genome reduction, some oomycetes have undergone substantial genome expansion (Raffaele & Kamoun, 2012). There is significant lifestyle diversity among these pathogens where some of them are obligatory biotrophs e.g. *Hyaloperonospora* sp.(Baxter et al., 2010); some are necrotrophs (e.g. many members of family Pythiaceae); some are hemi-biotrophs (e.g. many *Phytophthora* species); and some are saprophytes exhibiting significant environmental adaptability. The genome sizes of oomycete pathogens vary substantially, with the smallest one having 37 Mb (*Albugo laibachii*) and the largest one 240 Mb (*Phytophthora infestans*) (Pais et al., 2013).

Several oomycete pathogen genomes have been sequenced at different genome centers. However, most of the genome centers create their own databases for dissemination of data such as the Joint Genome Institute (JGI), Pythium Genome Database, Broad Institute etc. Some of these existing databases are on the verge of retirement and also do not have all the available oomycete genomes. For example, the Broad Institute’s resources recently closed. FungiDB hosts many fungal and oomycete genomes, but it is a very extensive resource, more appropriate for complete genomes having exhaustive functional annotation data. Eumicrobedb on the other hand is well suited for draft genomes that are still undergoing changes in terms of genome assembly and annotation. Changes made to a genome can be quickly and easily incorporated into EumicrobeDB. The data in this database have been integrated from different sources, so the nomenclature followed by the different centers had to be unified. We have adopted a standard system of nomenclature that is applicable to all the genomes. This system includes different assembly versions, annotations and the nomenclature of the features. The entire database package comprises of ~180,000 lines of code. The database resource is publicly available at www.eumicrobedb.org.

## Results and Discussion

Eumicrobedb has been significantly upgraded from its earlier version, VMD (VBI Microbial Database) in terms of functionality and content. Some of the advanced features are discussed below:

## Eumicrobedb runs on Genome Annotator Lite

EumicrobedbLite is an advanced version of VMD (VBI Microbial Database) (Tripathy et al., 2006), with major changes in its architecture and functionality. VMD was built on the Genome Unified Schema (GUS) that was based on an Oracle framework and had many interdependent bioperl modules for data integration and analysis. EumicrobedbLite on the other hand, is independent of proprietary software and external modules. It runs on a powerful genome analysis virtual machine - Genome Annotator Lite (GAL) developed at our lab at the Indian Institute of Chemical Biology, Kolkata, India (Panda et. al unpublished). GAL is a powerful yet lightweight virtual machine with most of the open source genome annotation tools embedded in it [manuscript under preparation]. In addition, the data parsing scripts in GAL do not require Bioperl, since inclusion of Bioperl makes installation of the package cumbersome. The workflow of GAL is illustrated in Figure 1. The entire process of uploading a completely annotated genome takes about 4 hours (for a genome with ~20,000 predicted genes). Alternatively, unannotated draft assemblies can be used as inputs that will be subsequently annotated, parsed and uploaded to the database by GAL; this process takes slightly longer, depending upon the genome size and the amount of analysis needed.

**Figure 1:**
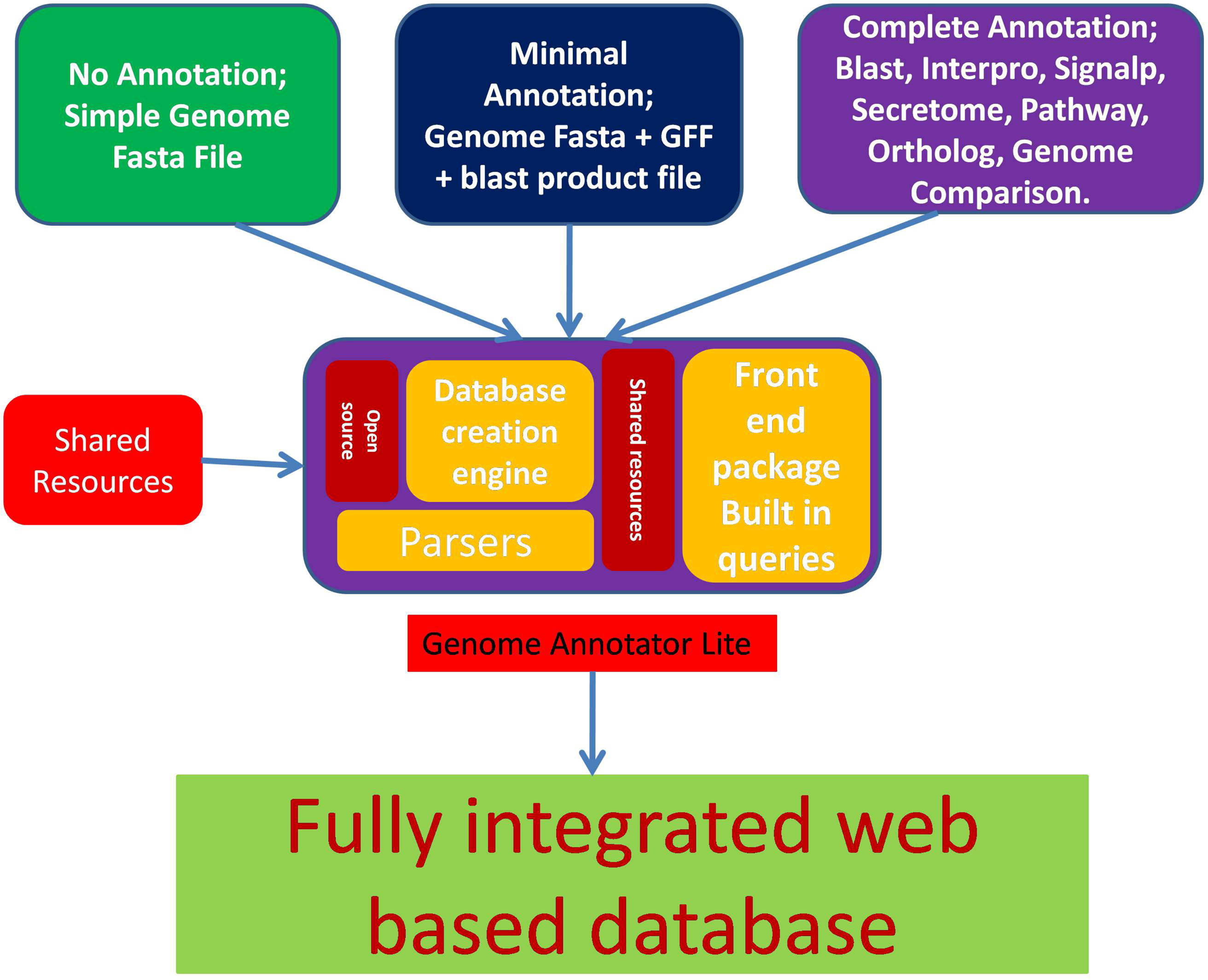
An overview of the GAL (Genome Annotator Lite) Workflow. Three different types of data can be provided as input to GAL: un-annotated, partially annotated and fully annotated. Depending upon the input type, GAL determines the type of analysis needing to be run on the data. GAL has the unique capability of creating a database schema if it is not present already. It can also download shared resources from public databases, parse the data and upload it into EuMicrobeDB.

Although genome sequencing technologies have been tremendously improved, annotating and visualizing them still remain a big challenge (Yandell & Ence, 2012). When a genome is sequenced for the first time, the first and foremost step is to clean the reads and assemble them and this is usually the most computationally intensive procedure. Currently, GAL does not handle genome assembly, since genome assembly can be extremely compute-intensive for many genomes. However, it takes care of the downstream data annotation. Users can submit data to GAL either as a draft assembly in a partially annotated form or as a fully annotated assembly. GAL will automatically recognize the data type and determine the type of analysis that is required. Currently, EumicrobedbLite contains analyzed genomes of 26 oomycete pathogens including one near complete genome of *Phytophthora sojae* (V5). Out of the 26 listed genomes in EumicrobedbLite, 21 genomes have been published while 5 are unpublished genomes that have some restrictions on use, namely *Phytophthora parasitica, Saprolegnia diclina, Aphanomyces astaci, Aphanomyces invadans* and *Phytophthora cinnamomi.* These 26 organisms are from different orders of the phylum Oomycota e.g. *Albuginales (Albugo sp.* - white rusts), *Peronosporales* (including plant pathogens such as *Phytophthora sp.* and downy mildews), *Pythiales (Pythium sp.*, water molds), *Saprolegniales (Saprolegnia sp.*, fresh water molds; many animal pathogens). Some organisms such as *Phytophthora sojae* have several different assembly versions and one of the earlier versions (version 1) is still widely used by researchers. So, we have included both genome versions of *Phytophthora sojae* (i.e. version 1 and version 5). The details of the number of scaffolds, genes, genome size, organism prefix etc. are available in the statistics link of the page [Supplementary Table1]. All the publicly available EST sequences of oomycete pathogens with their genome alignment data are also integrated into the database.

## EMBOSS Analysis package is integrated into the database

The sequence analysis package EMBOSS is a powerful tool comprising of 245 light weight sequence analysis programs (Rice et al., 2000). Out of these, we found 92 programs unsuitable or redundant in nature for web based applications. So, we have incorporated the remaining 153 useful programs from the EMBOSS package into the toolkit section of this database. Several packages extremely useful for sequence analysis have been integrated into the oomycete genomes present in the database, so that the users can select the genomes of interest through a drop down menu and perform the desired analysis on them. A few examples of these are listed in Table 1.

**Table 1:**
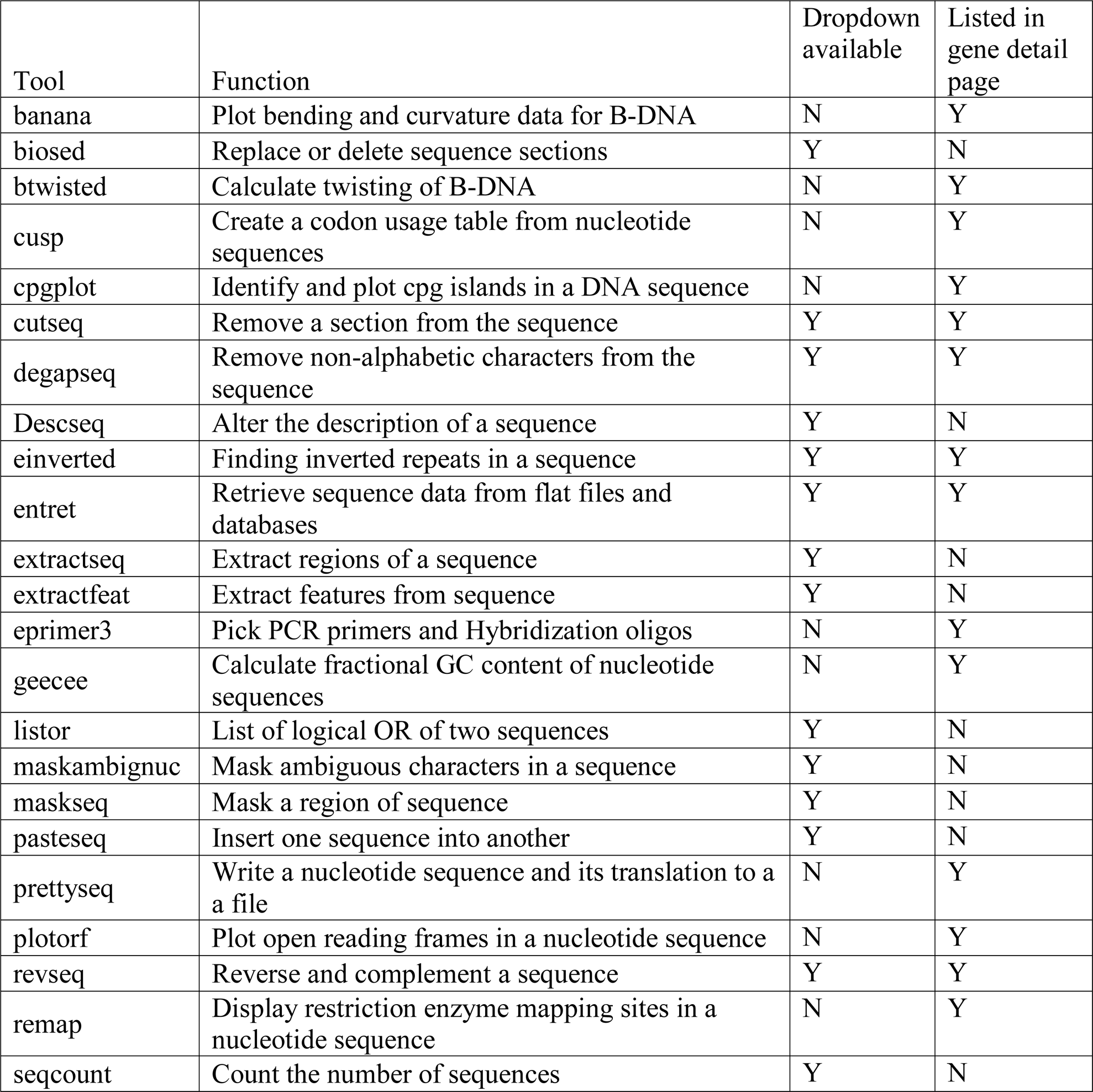
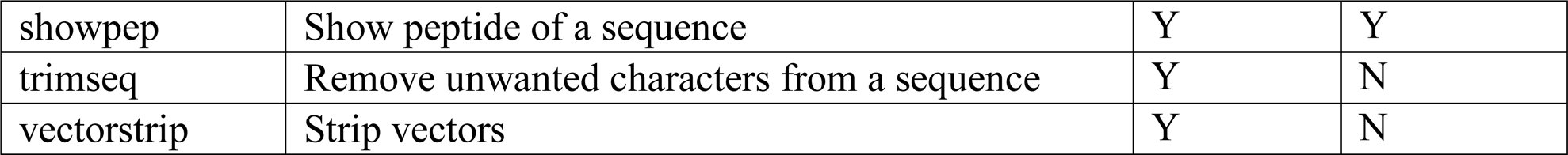
List of EMBOSS programs available as drop down menus built upon oomycete genomes that are listed in gene detail page.

Many sequence utility programs can be accessed from the gene detail page [Table 1]. These are linked to the gene models, so that with a single click users can run an analysis using the nucleotide or the protein sequence as automatic input. If the user chooses to run more than one analytical tool from this page, then the outputs are arranged in a tab separated menu. A “clear all” option is available to remove all analysis results from the page.

The analytical interfaces provided in Eumicrobedb provide a very simple and intuitive way to quickly run a variety of sequence analysis programs. The existing oomycete databases such as fungidb (Stajich et al., 2012), JGI’s Mycocosm (Grigoriev et al., 2014), the Pythium functional genomics database at Michigan State University (Hamilton et al., 2011) etc. disseminate precomputed genomics and comparative genomics data, but lack this feature for users to access a web-based analysis platform.

## Manually curated datasets for oomycete genomes are available in Eumicrobedb

In addition to the automated genome annotations, we have also carried out extensive semiautomated annotation and data curation of the oomycete genomes. One such curated data resource is the Core Ortholog dataset. The Core Ortholog dataset was obtained after comparing the entire proteomes of 12 representative members belonging to 4 different orders namely: *Albugo laibachii* Nc14 belonging to order Albuginales; *Phytophthora sojae* P6497, *Phytophthora ramorum* Pr102, *Phytophthora infestans* T30-4, *Phytophthora capsici*LT1534, *Phytophthora parasitica* INRA-310, *Hyaloperonospora arabidopsidis* Emoy2, *Plasmopara halstedii, Phytopythium vexans* DAOMBR484 (belonging to Peronosporales); *Pythium ultimum DAOMBR144 (V1*) (belonging to order Pythiales) and *Saprolegnia parasitica CBS223.65 (V1*), and *Aphanomyces invadans* (belonging to order Saprolegniales).

The Oomycete Molecular Genetics Network (OMGN at omgn.org) has conducted genome sequence jamborees for a number of newly released oomycete genome. In these jamborees community scientist members get together and manually annotate the data; the manual annotation effort often continues after the jamboree also. Over the last several years, we have collected this annotation data from community members and incorporated it into this database.

### a. Core Ortholog dataset for oomycete pathogens

Although many oomycete genomes are available, an organized effort to generate a core proteome was needed. We generated a comprehensive core proteome by choosing 12 representative oomycetes. Since all the genomes available currently are draft genomes, an attempt to include all 26 genomes for this study would have resulted in no core data set.

Initially we generated pairwise bidirectional best blast hits from these 12 proteomes (207,636 total protein sequences) followed by orthoMCL (Fischer et al., 2011), producing a set of core orthologs. We also did ortholog finding using multiparanoid (Alexeyenko et al., 2006) for comparison. The total numbers of core orthologs generated by multiparanoid and orthoMCL were comparable to each other i.e. 2862 and 2974 respectively. We used the core set generated by OrthoMCL since a number of clusters produced by multiparanoid were not very reliable. The clusters generated by OrthoMCL encompassed 44,777 protein sequences. The largest cluster has about 275 members (group 1) belonging to the ABC transporter superfamily [Supplementary Table 2; Supplementary Table 3; Supplementary Figure 1].

Out of the 2974 clusters generated using OrthoMCL, annotations of the individual clusters were obtained from the COG IDs of their members. Only about 1894 had COG annotations and the remaining 1080 are unique to this group of organisms. The greatest numbers of groups were annotated as Hypothetical Protein, Unnamed protein product, or conserved hypothetical protein. Among the annotated group members, the predominant classes of protein families were protein kinase [52 groups], ATP binding cassette [49 groups], transmembrane protein [49 groups], vacuolar associated proteins [31 groups], or serine protease family [31 groups] [Supplementary Figure1]. The ortholog clusters are available for search through the query page in eumicrobedb.org using two different options namely: query by cluster_ID or by annotation of clusters. Cluster information and the tree structure for a protein are also available on the protein’s gene detail page. A pre-computed cluster analysis for core orthologs is a very valuable resource for inferring the biological role and phylogeny of a protein sequence (Yang et al., 2015).

We generated an HMM profile of each of the orthologs cores using HMMER 3.1(Johnson et al., 2010). Then the database containing the protein sequences of the remaining 14 organisms (222,582 sequences) was searched against the HMM profiles using hmmsearch with a cutoff of 1e-05. A matrix was generated from the output with present calls as 1 and absent calls as 0. The distance between a pair of gene clusters in 2 genomes were calculated using the Jaccard distance method followed by single, complete and average clustering methods implemented in the Vegan Package in R (Scaria et al., 2015). Jaccard distance has been used widely to examine genome fluidity. A value close to zero means no difference between two genomes. We computed Jaccard distances between the set of 12 organisms as one group and the single genomes of the remaining 14 organisms as the other [Supplementary File 4; Figure 2]. *Phytophthora* taxon *totara* has diverged least from the cluster whereas most of the *Pythium sp.* have diverged the most. Some of these divergences may reflect variations in the quality of the respective assemblies and annotations. *Phytophthora sojae* (V1) records a very high distance (between a range of 0.04-0.05) compared with the others, indicating it is the least complete genome [Supplementary File 4].

**Figure 2:**
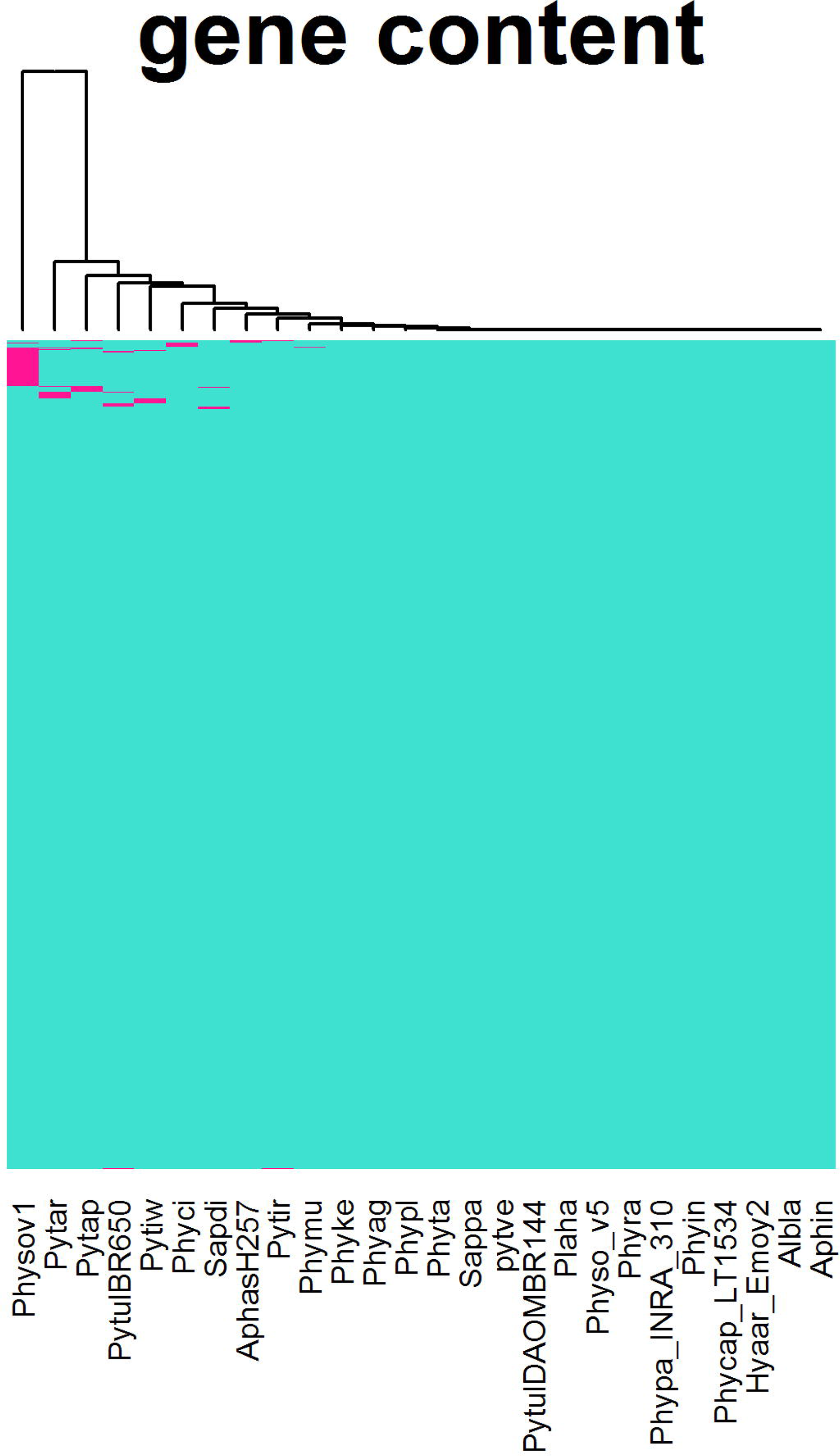
Heat map of the Jaccard distance between the core group and the remaining 14 organisms using single, complete and average linkage clustering using Vegan package in R.

### b. User annotated genes and effectors

Effectors are virulence proteins that enter host cells to promote infection. They undergo rapid evolution to evade detection by the host resistance machinery (Jiang et al., 2008). Since their rate of adaptation is very high, the sequences undergo rapid changes in composition. Gene predictors are therefore often unable to successfully predict these genes in a draft genome. Prediction of effectors requires the use of HMM searches and manual intervention in many cases. Most of the oomycete RxLR effectors have been curated by community users manually in conjunction with a variety of prediction strategies (Jiang et al., 2008) Haas et al., 2009). For those species with manually curated effector sets (*P. sojae, P. ramorum* and *Hyaloperonospora arabidopsidis*) we have replaced the electronically annotated effector sets with the manually curated gene models in this database version. Presently, there are about 125 curated RxLR effectors from *Hyaloperonospora arabidopsidis*; 370 curated RxLR effectors from *P. ramorum*; 396 curated RxLR effectors from *Phytophthora sojae* (V1) and 385 curated RxLR effectors from *Phytophthora sojae* (V5) [Table 2]. There are 1898 user-annotated non-effector gene models in the database. The user-predicted gene models as well as the gene models reviewed by the community reviewers are color coded in the browser (Figure 3B). Earlier versions of eumicrobedb (VMD) had a user annotation interface. However, users prefer to send bulk annotations in excel files rather than filling in gene details one by one in the web based forms.

**Figure 3:**
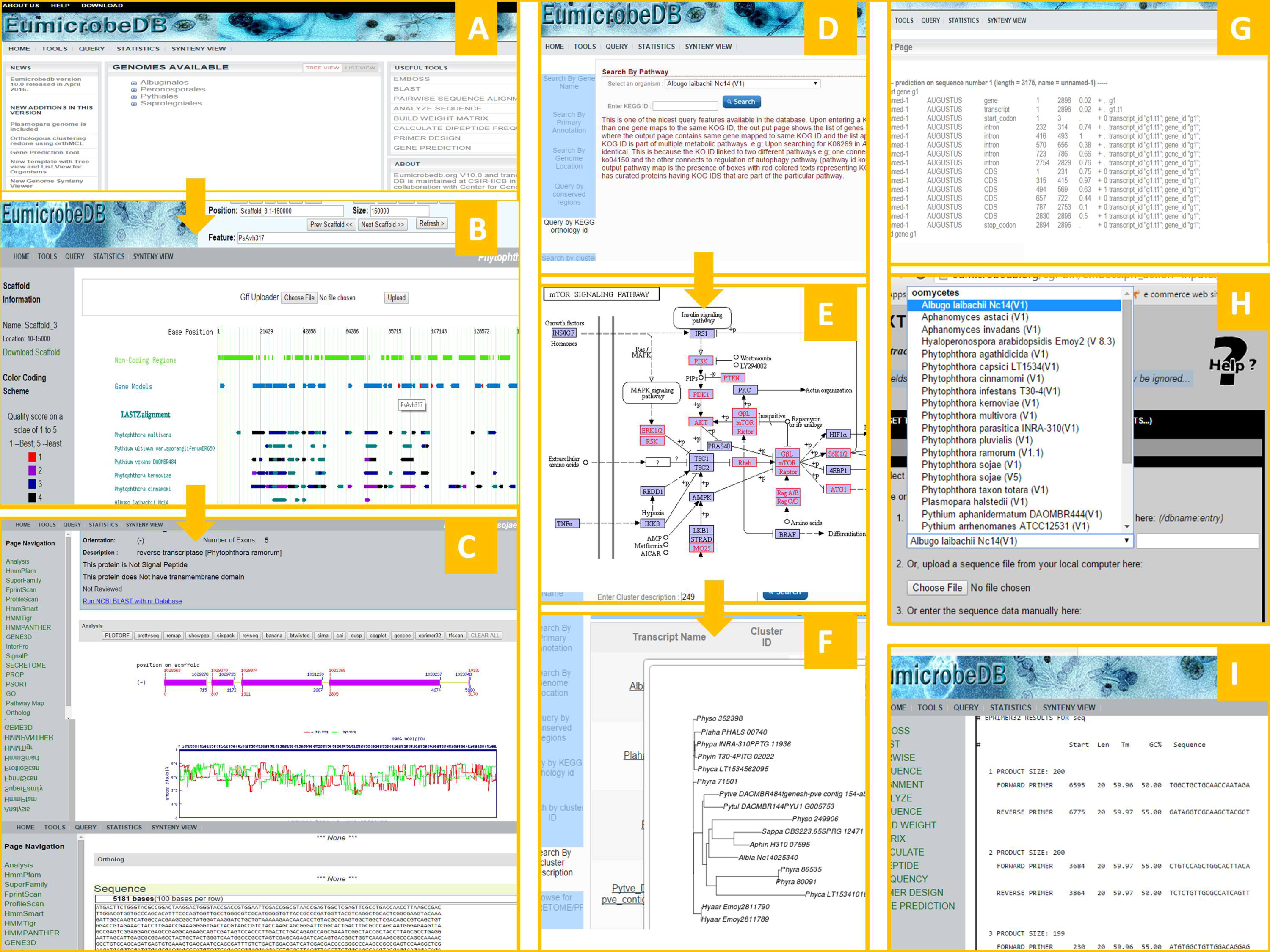
Graphical User Interface of Eumicrobedb. Screenshots are shown in each case. The home page (A) opens to a tree view list of the genomes present in Eumicrobedb. The genome browser page (B) opens showing the default scaffold with all available tracks. The tracks are clickable and clicking on a coding region opens to the gene detail page (C). Query items include KEGG ID (D), which leads to a pathway map (E) where the colored members are the KOGs present in the query organism for that pathway. The cluster query output (F) showing phylogenetic relationship between the genes from the same cluster. Gene prediction output (G), extract sequence with pull down genome menu (H) and output of primer design (I): All part of the new Toolkit menu.

**Table 2:** Number of curated RxLR effectors in Eumicrobedb

| Organism | Total |
| --- | --- |
| *Phytophthora sojae* P6497 (V1) | 396* |
| *Phytophthora sojae* P6497 (V5) | 385* |
| *Hyaloperonospora arabidopsidis* Emoy2 | 125*† |
| *Phytophthora ramorum* Pr102 (V1) | 370* |
| *Phytophthora cinnamomi* CBS 144.22 | 8 |
| *Phytophthora capsici* LT1534 | 159 |
| *Phytophthora infestans* T30-4 | 563‡ |

\* manually curated
† does not include 242 RxLR-like (RxLL) proteins judged to be poor quality effector candidates nor 22 crinkler genes with RxLR strings (RxLCRN genes).
‡ from Haas et al (2009)

So, we have taken off that feature in this version. Now the researchers can send us the gene related information through the ‘contact us’ page.

## Data visualization interface has many additional features

The data visualization interface of the database has five major components, namely: genome browser, gene detail page, genome synteny page, query page, and toolkit page. Several other accessory components such as statistics page, download page, and tutorial pages are also available. The gene detail page is the central part of the user interface where detailed annotation and analytical information is available for a gene. All the other pages eventually link to the gene detail page. A brief overview of the user interface is summarized in Figure 3.

### a. Synteny Viewer

The newly created synteny viewer is based on Genome Synteny Viewer - GSV (Revanna et al., 2011). Genome synteny was computed by running an all-versus-all comparison among all pairs of oomycetes genomes using Lastz (Harris, 2007). The user interface has been modified so that it is intuitive for new users. For example, users can choose to see the highly syntenic regions between a pair of organisms by just clicking on the ‘check synteny’ option. The scaffolds that display the most synteny between a pair of organisms will be listed on the page. Syntenic regions are displayed only if at least 10000 bases are syntenic and the insertions and deletions cover less than 5% of the matching length [Figure 4].

**Figure 4:**
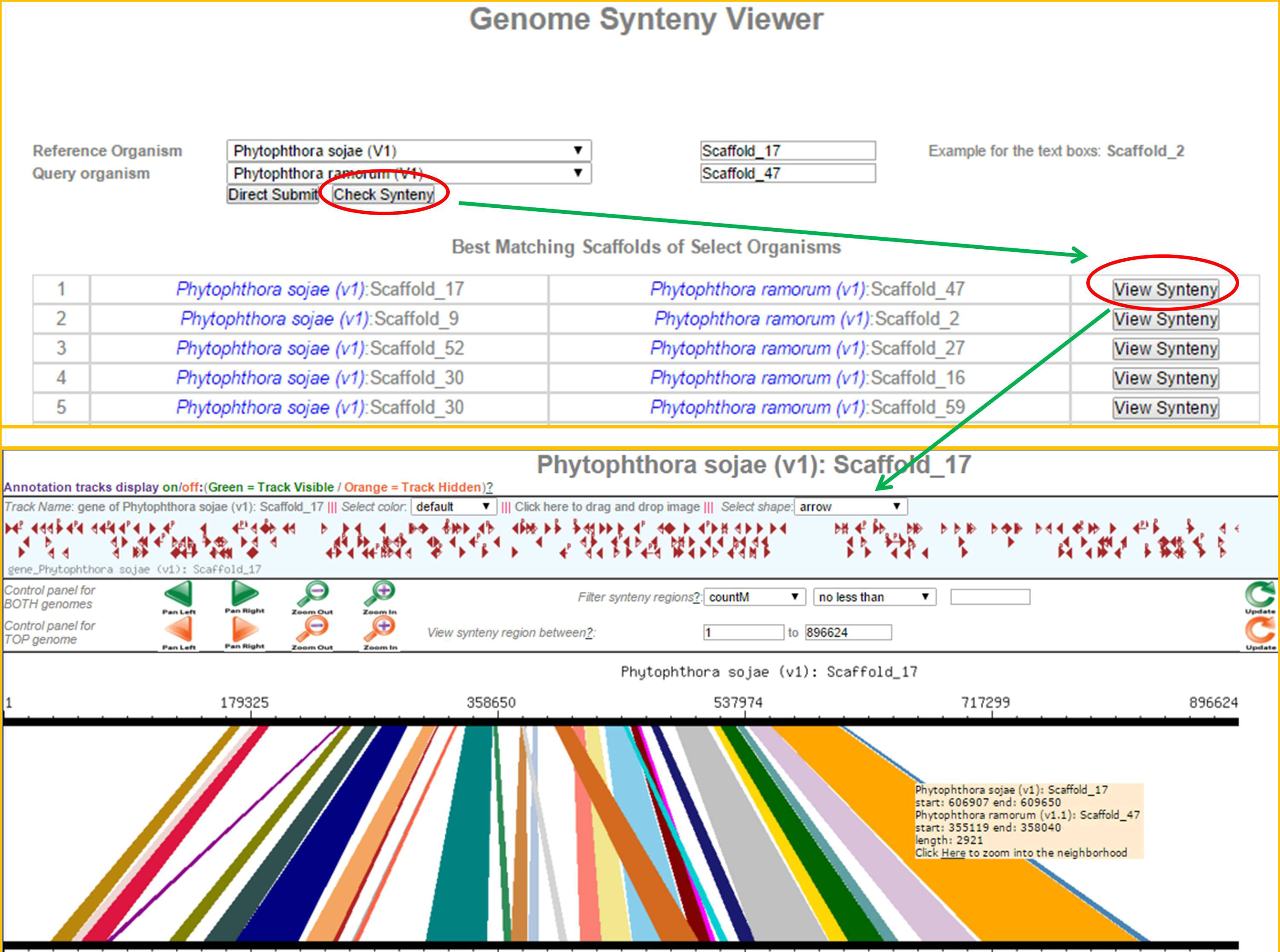
Scaffold synteny comparison. (A) Screenshot of the “check synteny option” between two specified scaffolds, which is one of two synteny query options (the other is querying the best syntenic regions for a particular scaffold - see Figure 4). A query is depicted between Scaffold_17 of *Phytophthora sojae* V1 and Scaffold_47 of *Phytophthora ramorum* V1. (B) All possible syntenic regions are listed for the user to choose to visualize synteny between a pair of scaffolds.

### b. Genome Browser

The genome browser serves as one of the entry points to this database from the main page. The organism list is arranged in a tree view format with taxonomic hierarchy e.g. orders-> Genus -> species->strains (if available); and in a list view format. On clicking an organism name, the browser page opens to the default scaffold page (largest scaffold) with default scaffold region (1-150,000). The uppermost green-colored track represents non-coding DNA sequences. This track is very useful when users are interested in retrieving upstream or downstream regions of a coding sequence.

Non-coding regions of the genomes are particularly interesting in the context of the ENCODE (2012) project. Natural selection plays a very important role in determining virulence of a pathogen and it may act on non-coding as well as coding regions of the genome (Rech et al., 2014). By offering the clickable non-coding track, researchers can quickly analyze the noncoding regions.

The next blue colored track identifies coding regions with introns and exons plotted as pointed rectangles indicating their orientation. The gene model is colored red when a community researcher has either reviewed or curated it. There are other feature tracks such as repeats and tRNA tracks available currently. We have added a new feature into the browser tracks, namely the conserved region track. The pre-computed comparative genomics regions between the genomes are quality-sorted and represented in 5 different colors on these tracks. The best quality conserved regions are ranked as 1 (color coded in red) and the least is ranked as 5 (please refer to the methods section for details about the scoring schemes). On ‘mouse-over’ on the conserved region tracks, the scaffold location of the conserved region pops up in a text box. This track is clickable and opens to a page containing the list of coding transcripts present in that region. Next to these tracks are the EST BLAT alignment tracks showing regions mapping between the reference genome and the assembled ESTs.

The genome browser and genome synteny viewer offer a quick and easy way to explore regions of a genome where genes are co-localized or where there is a repeat-mediated expansion of the genome [Figure 5; Figure 6].

**Figure 5:**
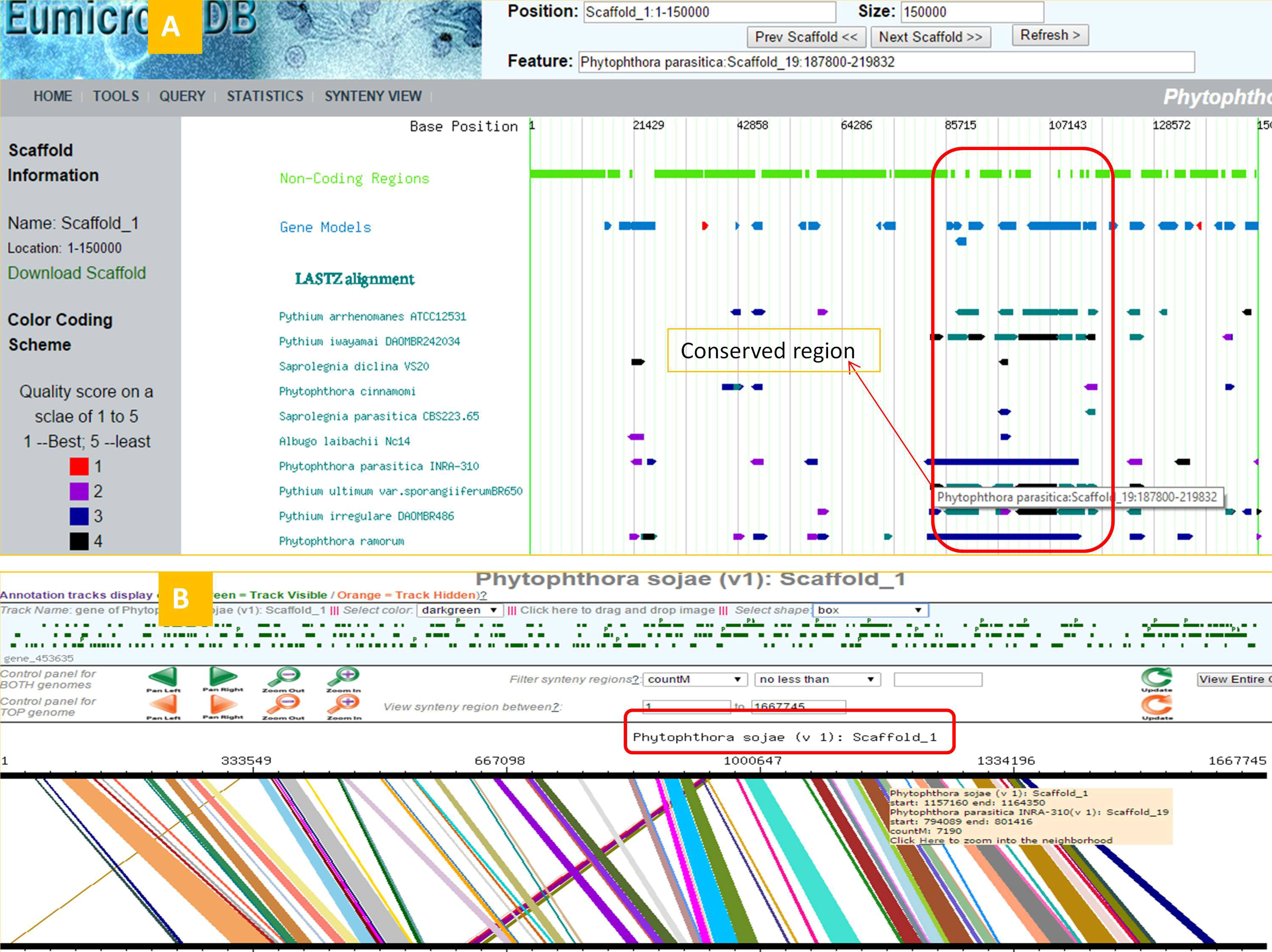
Scaffold genome-wide synteny query. (A) Screenshot of browser page showing synteny between scaffold_19 of *P. parasitica* with conserved regions of other oomycete genomes. *P. sojae* scaffold_1 shows significant synteny with scaffold_19 of *P. parasitica.* (B).

**Figure 6:**
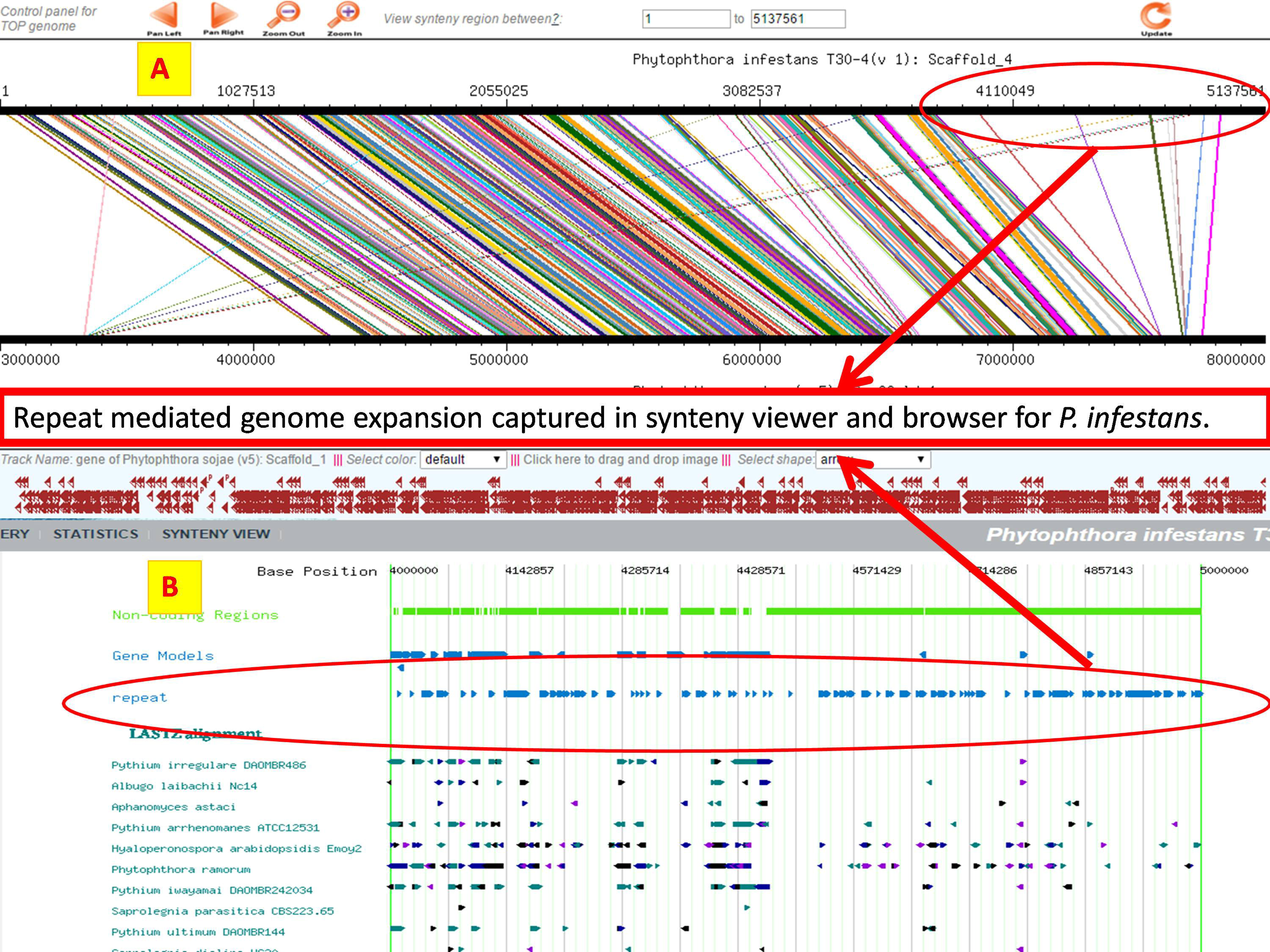
Repeat mediated genome expansion in *Phytophthora infestans* illustrated with the synteny viewer and browser. (A) Synteny view of *P. infestans* scaffold_4 aligned with *Phytophthora sojae* V5 scaffold_1 (B) Browser view of *P. infestans* showing region of scaffold_4:4000000-5000000.

Users can now upload data in GFF format into the browser for quick visualization [Template file provided in download page at [http://www.eumicrobedb.org/uploadable_gff/]. The coding tracks of the user-uploaded gff tracks are clickable.

### C. Query Page

The updated query page is frame-based, where different kinds of queries can be carried out by clicking on the left menu item. Each query result is then stored in the browser cache for easy data browsing. The query options include: query by gene_id, query by primary and secondary annotation, query by GO annotation, query by cluster_ID, query by cluster annotation, EST query, synteny query, query by KEGG orthology id etc. Bulk data download for the secretomeP package (PROP, SIGNALP, TMHMM, PSORT) is available via this page. Query outputs for genes open into the relevant gene detail page. Query results that contain multiple transcripts or genes open as a list where each item is linked to the relevant gene detail page. “Query by genome location” retrieves a list of features that occur within the queried region, together with links to the relevant gene pages. “Query by conserved region” is a new feature that is similar to “query by genome location”, except that the output is a set of features contained in regions of other genomes that are conserved with the query region. “Query by KEGG Orthology” results in a list of the genes tagged to the KEGG ID query. Upon clicking the KEGG IDs, the pathway image maps appear with colored EC numbers. These colored EC numbers indicate members of that pathway present in the reference genome.

### d. Gene detail page

The gene detail page or main annotation page contains detailed information about a gene. This page has a summary header containing brief information on the gene/transcript. Since this page contains a long list of information, quick links to different features are provided in the left panel. Loglikelihood (McLachlan et al., 1984) and Fickett statistics (Fickett, 1982) plots of the genes are computed on-the-fly using pre-computed codon-usage values. These plots help validate the correctness of the predicted gene model. The gene model plot on the top is clickable and the page leads to the translated CDS and nucleotide sequences for the gene. A new analytical feature is incorporated in the top panel that runs some of the sequence analysis programs chosen from

EMBOSS. The users can click on the tools and the gene sequence in the page will be used as the input for the program and output will be displayed in the same page.

### e. Toolkit Page

The toolkit page is the sequence analysis interface of eumicrobedb with many useful open source as well as in-house tools for sequence analysis. Blast, Pairwise sequence comparison, and the EMBOSS interface are a few of the most useful packages that are part of this suite. From the EMBOSS package, 150 sequence analysis programs are integrated with the 26 oomycete genome sequences. The inputs to many of these programs can be selected from the drop down menu box and analysis can be carried out directly. By integrating open source sequence analysis packages that normally exist as stand-alone packages, eumicrobedb provides a convenience that is invaluable for biologists.

Another very useful feature added to the toolkit is the integration of the gene prediction software Augustus (Stanke et al., 2008). We have refined training datasets for each of the 26 oomycete genomes and users can choose the training dataset of their choice for predicting coding regions from an unknown stretch. This is one of the most useful features for the research community.

## Experimental Procedure

Genomes were downloaded from genome center web sites. The sources and origins of the genomes are listed on the Statistics page of the eumicrobedb web site and EST datasets were downloaded from Genbank.

## Data Processing

Genome sequence files and gene names have been renamed with genus, species and strain prefixes for uniformity. For genomes such as *P. sojae* where more than one assembly version available, we have appended the assembly version to the genus_species_strain prefix. For unification of scaffold nomenclature, we have size-sorted scaffolds and numbered them in descending order from larger to smaller, thus each largest scaffold is named as Scaffold_1. For some organisms such as *Hyaloperonospora arabidopsidis* and *Phytophthora infestans* where the scaffolds already followed this rule, we kept the old name. A map file is provided at www.eumicrobedb.org/ForEMBOSS/ for comparing the old names with the new names. Genes and genome prefixes are listed on the Statistics page.

We analyzed and annotated 406,500 protein coding genes from these 26 oomycete organisms. Blastx (Altschul et al., 1990) against NCBI’s nr database was used to assign putative primary annotations to the genes. We ran interproscan (Zdobnov & Apweiler, 2001) annotation for predicting domains and GO features. Pathway and KOG prediction was done using KAAS annotation server from KEGG (Moriya et al., 2007). Additionally, we created an entire secretome repertoire using SignalP 3.0 (Petersen et al., 2011), and the secretomeP (Bendtsen et al., 2004) package; the latter includes Prop (Duckert et al., 2004) (Prediction of Proprotein convertase sites), Psort (Horton & Nakai, 1997) (prediction of protein sub-cellular locations), and TMHMM (Krogh et al., 2001).

For creating a core proteome of the oomycetes, we used 12 oomycete organisms as described in the core ortholog generation section of the Results and Discussion. Clusters of orthologous proteins were extracted from 12 organisms using OrthoMCL version 2.0.9 using default parameters. A total of 207,636 proteins were clustered into 22,592 groups. Core orthologs were defined as the 2,974 ortholog groups that were present in all 12 organisms. In order to detect core orthologs in the remaining 14 organisms (out of the total 26), we did a profile search based on HMMER. Each of the 2974 clusters contained multiple proteins and were separated into individual clusters. Profiles were built from each of the 2974 clusters and an HMMSearch was carried out against these 2974 profiles for all the 14 other organisms that were not part of the cluster building. A matrix of 1s and 0s was constructed for the 25 organisms consisting of 2974 rows. This matrix was then converted into a distance matrix using Jaccard Distance implemented in the Vegan package in R. Hierarchical clustering was performed on the resulting distance matrix using “single”, “complete” and “average” linkage clustering as implemented in Vegan. Heatmaps were drawn to visualize the presence/absence with the stats package in R. Annotations of the clusters were manually edited to fit into a broader category before uploading into tagcrowd.com for generation of word clouds.

We built phylogenetic relationships among these proteins with ClustalW (Thompson et al., 2002) and MEGA (Tamura et al., 2013). The Multiple sequence alignment and the tree features are available in gene detail page.

We have included secretomeP analysis for predicting non-signal peptide secretory proteins Whole genome synteny analysis was done by running all vs. all whole genome comparisons using LastZ (Harris, 2007). EST data sets were cleaned using in-house scripts, clustered and assembled using TGICL (Pertea et al., 2003). EST contig alignment to genome assemblies was done using BLAT (Kent, 2002).

## Comparative Genomics Module

We aligned all the existing 26 genomes against each other (624 runs) using Lastz, a package that handles pairwise sequence alignments (Harris, 2007). We performed chained, gapped alignments with the default mismatch count (< 50 mismatches) for Lastz over windows of 1000 bases. The alignments were further filtered into 5 categories with the best being 1 and the least match being 5. The best matches have a matching region of over 10,000 bases with < 5% mismatches and gaps. Second best matches have >1,000 bases and <10,000 bases matching region with < 5% mismatches and gaps. The third category is for matching regions over 1000 bases with mismatches and gaps > 5% and < 10%. The fourth category is for matching regions over 1000 bases with mismatches > 10% and < 15%. The remainder are category 5.

## Future Directions

Many new oomycetes genomes are being sequenced at several genome centers. We are on our way to collecting the publicly available genomes into this database in the next release.

## Acknowledgement

### Availability

Eumicrobedb.org is publicly available at www.eumicrobedb.org.

### Authors Contribution

ST and BMT designed the project and wrote the manuscript; ST, MMC, DS, AD, GPM and SD analyzed the data; AP, AG and ST uploaded the data; AP designed and created the front end, WWY and BMT provided curated lists of RxLR effectors.

### Acknowledgement

Financial support from DBT-RLS fellowship, Department of Biotechnology and CSIR-Genesis, Govt. of India to ST is gratefully acknowledged. This work was supported in part by grants to B.M.T. from the USDA National Institute of Food and Agriculture, #200735600-18530 and #2011-68004-30104 and from the US National Science Foundation, #MCB-0731969.

## Supplementary File Legends

Supplementary Figure1 : Word cloud for 100 most frequent words in the annotation file for 2974 core groups computed using OrthoMCL with 12 representative members.

Supplementary File1: Gene Statistics of all the members present in Eumicrobedb.

Supplementary File2: 2974 clusters generated with OrthoMCL with their COG IDs, annotations.

Supplementary File3: Members of each of the 2974 clusters with organism ids, gene_ids, cluster_id and annotation.

Supplementary File4: Distance matrix showing Jaccard distance between the cluster of 12 organisms with other members.

## References

Adhikari BN, Hamilton JP, Zerillo MM, Tisserat N, Levesque CA, Buell CR, 2013. Comparative genomics reveals insight into virulence strategies of plant pathogenic oomycetes. PloS one 8, e75072.

Alexeyenko A, Tamas I, Liu G, Sonnhammer EL, 2006. Automatic clustering of orthologs and inparalogs shared by multiple proteomes. Bioinformatics 22, e9–15.

Altschul SF, Gish W, Miller W, Myers EW, Lipman DJ, 1990. Basic local alignment search tool. J Mol Biol 215, 403–10.

Anonymous, 2012. An integrated encyclopedia of DNA elements in the human genome. Nature 489, 5774.

Baxter L, Tripathy S, Ishaque N, et al., 2010. Signatures of adaptation to obligate biotrophy in the Hyaloperonospora arabidopsidis genome. Science 330, 1549–51.

Bendtsen JD, Nielsen H, Von Heijne G, Brunak S, 2004. Improved prediction of signal peptides: SignalP 3.0. J Mol Biol 340, 783–95.

Dong S, Stam R, Cano LM, et al., 2014. Effector specialization in a lineage of the Irish potato famine pathogen. Science 343, 552–5.

Duckert P, Brunak S, Blom N, 2004. Prediction of proprotein convertase cleavage sites. Protein Eng Des Sel 17, 107–12.

Fickett JW, 1982. Recognition of protein coding regions in DNA sequences. Nucleic Acids Res 10, 5303–18.

Fischer S, Brunk BP, Chen F, et al., 2011. Using OrthoMCL to assign proteins to OrthoMCL-DB groups or to cluster proteomes into new ortholog groups. Curr Protoc Bioinformatics Chapter 6, Unit 6 12 1–9.

Grigoriev IV, Nikitin R, Haridas S, et al., 2014. MycoCosm portal: gearing up for 1000 fungal genomes. Nucleic Acids Res 42, D699–704.

Hamilton JP, Neeno-Eckwall EC, Adhikari BN, et al., 2011. The Comprehensive Phytopathogen Genomics Resource: a web-based resource for data-mining plant pathogen genomes. Database (Oxford) 2011, bar053.

Harris RS, 2007. Improved pairwise alignment of genomic DNA. Ph.D Thesis, The Pennsylvania State University.

Horton P, Nakai K, 1997. Better prediction of protein cellular localization sites with the k nearest neighbors classifier. Proc Int Conf Intell Syst Mol Biol 5, 147–52.

Jiang RH, Tripathy S, Govers F, Tyler BM, 2008. RXLR effector reservoir in two Phytophthora species is dominated by a single rapidly evolving superfamily with more than 700 members. Proc Natl Acad Sci U S A 105, 4874–9.

Johnson LS, Eddy SR Portugaly E, 2010. Hidden Markov model speed heuristic and iterative HMM search procedure. BMC Bioinformatics 11, 431.

Kent WJ, 2002. BLAT–the BLAST-like alignment tool. Genome Res 12, 656–64.

Krogh A, Larsson B, Von Heijne G, Sonnhammer EL, 2001. Predicting transmembrane protein topology with a hidden Markov model: application to complete genomes. J Mol Biol 305, 567–80.

Mclachlan AD, Staden R, Boswell DR, 1984. A method for measuring the non-random bias of a codon usage table. Nucleic Acids Res 12, 9567–75.

Moriya Y, Itoh M, Okuda S, Yoshizawa AC, Kanehisa M, 2007. KAAS: an automatic genome annotation and pathway reconstruction server. Nucleic Acids Res 35, W182–5.

Pais M, Win J, Yoshida K, et al., 2013. From pathogen genomes to host plant processes: the power of plant parasitic oomycetes. Genome Biol 14, 211.

Pennisi E, 2010. Armed and dangerous. Science 327, 804–5.

Pertea G, Huang X, Liang F, et al., 2003. TIGR Gene Indices clustering tools (TGICL): a software system for fast clustering of large EST datasets. Bioinformatics 19, 651–2.

Petersen TN, Brunak S, Von Heijne G, Nielsen H, 2011. SignalP 4.0: discriminating signal peptides from transmembrane regions. Nat Methods 8, 785–6.

Raffaele S, Kamoun S, 2012. Genome evolution in filamentous plant pathogens: why bigger can be better. Nat Rev Microbiol 10, 417–30.

Rech GE, Sanz-Martín JM, Anisimova M, Sukno SA, Thon MR, 2014. Natural Selection on Coding and Noncoding DNA Sequences Is Associated with Virulence Genes in a Plant Pathogenic Fungus. Genome biology and evolution 6, 2368–79.

Revanna KV, Chiu CC, Bierschank E, Dong Q, 2011. GSV: a web-based genome synteny viewer for customized data. BMC Bioinformatics 12, 316.

Rice P, Longden I, Bleasby A, 2000. EMBOSS: the European Molecular Biology Open Software Suite. Trends Genet 16, 276–7.

Scaria J, Suzuki H, Ptak CP, et al., 2015. Comparative genomic and phenomic analysis of Clostridium difficile and Clostridium sordellii, two related pathogens with differing host tissue preference. BMC genomics 16, 1–16.

Stajich JE, Harris T, Brunk BP, et al., 2012. FungiDB: an integrated functional genomics database for fungi. Nucleic Acids Res 40, D675–81.

Stanke M, Diekhans M, Baertsch R Haussler D, 2008. Using native and syntenically mapped cDNA alignments to improve de novo gene finding. Bioinformatics 24, 637–44.

Tamura K, Stecher G, Peterson D, Filipski A, Kumar S, 2013. MEGA6: Molecular Evolutionary Genetics Analysis version 6.0. Mol Biol Evol 30, 2725–9.

Thompson JD, Gibson TJ, Higgins DG, 2002. Multiple sequence alignment using ClustalW and ClustalX. Curr Protoc Bioinformatics Chapter 2, Unit 2 3.

Tripathy S, Pandey VN, Fang B, Salas F, Tyler BM, 2006. VMD: a community annotation database for oomycetes and microbial genomes. Nucleic Acids Res 34, D379–81.

Tyler BM, Tripathy S, Zhang X, et al., 2006. Phytophthora genome sequences uncover evolutionary origins and mechanisms of pathogenesis. Science 313, 1261–6.

Yandell M, Ence D, 2012. A beginner’s guide to eukaryotic genome annotation. Nat Rev Genet 13, 329-42.

Yang L, Tan J, O’brien EJ, et al., 2015. Systems biology definition of the core proteome of metabolism and expression is consistent with high-throughput data. Proc Natl Acad Sci U S A 112, 10810–5.

Zdobnov EM, Apweiler R, 2001. InterProScan–an integration platform for the signature-recognition methods in InterPro. Bioinformatics 17, 847–8.

